# Discriminating Mechanisms of Heart-Voice Coupling

**DOI:** 10.64898/2025.12.01.691544

**Authors:** Marijn S.J. Hafkamp, Raphael Werner, Linda Drijvers, Luc Selen, Wim Pouw

## Abstract

In a landmark paper in *JASA*, Orlikoff & Baken (1989a) demonstrated that the heartbeat carries through acoustic features of the human voice. Heart-voice coupling questions how vocal control is functionally regulated and has many potential engineering applications to extract heart rate from the human voice. However, laboratory phonology and bioacoustics seem to have forgotten about the phenomenon, such that its physiological underpinnings have been left unexplained. The current contribution showed that vocal acoustics contain signatures of the heartbeat as originally reported by Orlikoff & Baken (1989a). More specifically, the steady-state vowel vocalizations of 17 participants were analysed and a significant relationship between the vocal acoustic envelope and the cardiac activity (ECG) was found. Furthermore, two empirical tests were conducted to discriminate between vocal fold (vascularization) and pulmonary (heart-lung interaction) mechanisms of heart-voice coupling. In addition to a stable heart-voice coupling, a significant heart-expiration coupling during conditions of silent expiration was also found. Since the vocal folds are abducted during expiration, this result demonstrated that pulmonary mechanisms that modulate expiratory flow must be involved in heart-voice coupling. It is speculated that the heart’s rhythm may be a weakly coupled pattern generator that possibly shaped the evolution of vocal and speech production.

## I. INTRODUCTION

That pulses of the heart carry through the human voice is not a well-known phenomenon. It is absent from standard textbooks on the physiology of vocalization and speech (e.g., Seikel et al., 2015) and it is not an active topic of study in laboratory phonology, psycholinguistics, nor in comparative psychobiology. Yet, the existence of heart-voice coupling was already reported in 1989 in *The Journal of the Acoustic Society of America* (Orlikoff & Baken, 1989a). In a landmark study, Orlikoff and Baken showed that the fundamental frequency (*f*_0_) of a sustained vowel vocalization was systematically modulated by the beating of the heart. More precisely, the heartbeat accounted for about 5% of the variance in the *f*_0_, also known as vocal jitter (Orlikoff & Baken, 1989b). Follow-up studies replicated this result (Orlikoff, 1990a) and demonstrated that the (acoustic) amplitude of the vocalization was also affected by the cardiac activity. On average, the heartbeat explained 12% of the amplitude variance, the vocal shimmer, during sustained and steady-state vowel vocalization (Orlikoff, 1990b). Together, these findings suggest an intriguing connection between the cardiovascular system and the vocal system, which raises the question of how vocal control is maintained and has evolved under heart beat perturbations. In this paper, we replicate the phenomenon and investigate its underlying mechanisms.

The coupling of heart and voice has many potential applications. In medical fields, for instance, it can be used to develop non-contact mobile heart rate (HR) monitors from vocalization acoustics or from real time speech signals. This can support early recognition of cardiac arrest (Milton & Monsely, 2018). In psychology, heart-voice coupling can be used to derive cardiac correlates of emotional episodes (James, 2015; Smith et al., 2017). Likewise, the field of bioacoustics could potentially extract HR from wild animal call recordings. Though none of these technologies have been developed yet, such applications could monitor the health of wild animals, and most presciently they could benefit welfare monitoring of commercial farm animals (e.g., Reza et al., 2025). Interestingly, there is evidence for a mechanical interaction of the heart impacting the lungs in pigs, leaving a notable 15 ml lung volume of expiratory drive when the heart fills during the late diastole phase (Lichtwarck-Aschoff et al., 2004). In the technological domain, heart-voice coupling can also be used to help recognize deep fake videos, next to visual correlates of HR in for example the face coloration (Boccignone et al., 2022; Gyanchandani et al., 2024). If the HR extracted from the vocal acoustics in a video is not within biological boundaries, the audiovisual recording may be flagged as potentially fake.

A deeper understanding of heart-voice coupling may be of further interest for behavioral and neuroscientific disciplines. In recent decades, it has become increasingly clear that human behavior is not only regulated by our interaction with the external environment, but also by the physiological cycles within our own bodies (Duschek et al., 2013). To give just one example, the vocalization of infants appears to be timed to the rise and fall of the heart rate (Borjon et al., 2024), suggesting that the beating of the heart may serve as an initial scaffolding for learning speech or language. As such, a field like developmental psychology could benefit as much from knowledge about heart-voice coupling as more technologically oriented fields can.

In concordance with the potential impact of heart rate detection from vocal acoustics, the past few decades have witnessed the emergence of various algorithms to extract HR from voice recordings. Some of these algorithms have used steady-state vowel vocalizations (Mesleh et al., 2012; Sakai, 2015), in alignment with the original work of Orlikoff & Baken (1989a)^1^. Other algorithms, however, employed machine learning to produce predictions from complex and dynamic speech signals. James (2015) used voice recordings of sentences expressing an emotion (e.g., anger or joy), while Smith et al. (2017) based their predictions on voice recordings of participants interacting with a customer service that was set up to evoke negative responses. Typically, these studies extract multiple acoustic features from the speech signal, such as Mel Frequency Cepstral Coefficients (James, 2015; Milton & Monsely, 2018). Subsequently, they employ a host of available techniques to extract the HR from these features, ranging from naive Bayes classifiers (James, 2015) to linear support vector regressions (Schuller et al., 2013; Milton & Monsely, 2018), and from random forest modeling to LASSO (Least Absolute Shrinkage and Selection Operator) regressions (Smith et al., 2017). This results in prediction errors from 8 beats per minute (bpm, Schuller et al., 2013) to 13 bpm (Milton & Monsely, 2018), which is reasonable, but less accurate than the HR predictions based on simple vowel vocalizations (e.g., 4 bpm in Sakai, 2015).

The data-driven extraction of the heart rate from complex speech signals is an impressive achievement. But such technological progress will stifle if it is not accompanied with a deeper understanding of the mechanisms underlying heart-voice coupling. Furthermore, how humans maintain control of their voice under sustained cardiac perturbation requires understanding of the mechanisms of heart-voice coupling. Despite this scientific and technological relevance, research into the physiological or neural mechanisms behind the coupling has been near absent in the three decades following the work of Orlikoff and Baken (1989a). With a few exceptions (Mesleh et al., 2012; Sakai, 2015; Poleshenkov & Basov, 2020), most researchers who engaged with the topic did not formulate any hypotheses regarding the mechanisms of heart-voice coupling, nor regarding its impact on the control of speech. And even when such hypotheses were formulated, no attempts were made to falsify them in an experiment. A possible reason for the neglect is that data-driven engineering approaches are generally not concerned with the ‘how’ nor with the ‘why’ of a biological phenomenon. These questions are more appropriate for fields like experimental biology, laboratory phonology, or bioacoustics. Yet unfortunately, those very fields seem to have left heart-voice coupling unexplained, if not to have forgotten about the phenomenon altogether. Therefore, the current study aims to readdress heart-voice coupling as a topic of study for understanding vocalization and speech. Taking the work of Orlikoff (and Baken) as our point of departure, we test a series of hypotheses about the mechanisms behind heart-voice coupling.

### A. Potential mechanisms of heart-voice coupling

The acoustic features of the human voice are predominantly generated by the vibration of the vocal folds. The frequency of these vibrations relates to the perceived pitch of the voice, while their amplitude of motion determines to a large part its acoustic amplitude. Therefore, it is not surprising that the vocal folds are often designated as the “source” of heart-voice coupling (Orlikoff, 1989; Sakai 2015; Poleshenkov & Basov, 2020). Orlikoff (1989) suggested that the vibrational qualities of the vocal folds may be affected by the vascularization of the glottis. He speculated that the systolic increase in blood pressure may increase the stiffness of the folds, explaining the periodic rise in *f*_0_ at the beating of the heart (Orlikoff & Baken, 1989a). At the same time, they acknowledged that this effect might be counteracted by the increase in vocal fold mass during the systole, which in turn lowers the frequency of the fold vibrations. Therefore, it is also possible that the cardiovascular system affects vocal fold behaviour more indirectly, for example through the vascularization of the laryngeal musculature (Orlikoff, 1989; Mesleh et al., 2012). Laryngeal muscles, such as the thyroarytenoid, contract and increase volume during the systole. This narrows the glottis and thereby reduces the glottal closure time, increasing the fundamental frequency of the voice. These two mechanisms, to which we shall jointly refer as *vocal fold mechanisms*, have never been contrasted.

As mentioned before, Orlikoff (1990a, 1990b) found that the heartbeat not only affected the fundamental frequency of the voice, but also its acoustic amplitude. Although the acoustic amplitude is indeed partly determined by the vibrational qualities of the vocal folds, its prime determinant is the subglottal or pulmonary pressure (Finnegan et al., 2000). This suggests that the heartbeat may (also) manifest itself in the acoustic amplitude by causing fluctuations in the pulmonary pressure^2^. For instance, fluctuations in the blood pressure of the aorta, pulmonary artery, and lung vessels may affect the rate of the air flow from the lungs (Poleshenkov & Basov, 2020), causing the subglottal pressure to rise and fall with the beating of the heart. Additionally, the heartbeat may have a mechanical impact on the lungs^3^ (Orlikoff, 1989; Orlikoff, 1990b). Because the heart is connected to the lungs via the pericardium, the pumping of the heart could cause fluctuations in the alveolar pressure with the heartbeat. Note, however, that these fluctuations may have a different phasing than those caused by the flow of blood through the lungs. Whereas contraction of the heart would increase the blood pressure, it would simultaneously decrease the mechanical pressure, because the size of the heart reduces during contraction. Which of these *pulmonary mechanisms* exerts the largest influence on the vocal system has been left to speculation. In fact, empirical evidence for any of the hypothesized mechanisms –vocal fold or pulmonary– is currently lacking.

In sum, there is no consensus in the literature on the physiological mechanisms of heart-voice coupling. Most papers seem to regard the vascularization of the vocal folds and laryngeal musculature as the most likely cause of the heart-voice coupling, yet the significant impact of the heartbeat on acoustic amplitude also suggests a role for subglottal interactions between the heart and the lungs. The current study is designed discriminate between these two mechanisms.

### B. Current study

To find signatures of the heartbeat in the voice, we investigated the acoustic amplitude of sustained vowel vocalizations. More specifically, we studied the squared magnitude coherence between the acoustic signal and the cardiac activity of vocalizing participants. An above-chance coherence would replicate the findings of Orlikoff (1990a, 1990b) and indicate a coupling of voice and heart. However, it would not yet discriminate between vocal fold and pulmonary mechanisms in the heart-voice coupling. To contrast these, we designed two additional empirical tests.

As a first test, we contrasted heart-voice coupling at the beginning and at the end of the vocalization, under the hypothesis that this coupling should weaken if it has a pulmonary origin^4^.

Over the course of a vocalization, the lungs reduce in air volume and the expiratory drive of the elastic recoil of the breathing system diminishes. Without countermeasures of recruiting additional respiratory muscles (e.g., Bouhuys et al., 1966; see also Ohala, 1990) this may lead to gradual decline in the pulmonary pressure and a decrease in the acoustic amplitude. If the heart-voice coupling has a pulmonary cause, one expects the manifestation of the heartbeat in the acoustic amplitude to decrease when the lungs are deflating. As such, we compared the coherence between the acoustic signal and the cardiac activity in the first half of the vocalization to the coherence in the second half. Higher coherence in the first half compared to the second half would provide evidence for pulmonary mechanisms underlying heart-voice coupling. Conversely, finding no significant difference would suggest that heart-voice coupling is not dependent on the deflation of the lungs, and potentially not dependent on the lungs at all.

Another way to discriminate vocal fold and pulmonary mechanisms is to investigate heart-voice coupling during silent expiration, without speech. In expiration, the vocal folds are abducted. This excludes mechanisms such as the vascularization of the glottis as a physiological cause for heart-voice coupling, because abducted vocal folds cannot vibrate during expiration. Thus, if the heartbeat can still be found in the amplitude of a silent expiration, then pulmonary mechanisms *must* be involved in heart-voice coupling, which would be in line with findings that the acoustic amplitude is a key acoustic correlate of the heart pulse. In the same way that we studied vocalization, we computed the coherence between the amplitude signal and the cardiac activity during expiration conditions. Finding above chance coherence values would provide evidence for pulmonary mechanisms. Conversely, finding no coherence would firmly exclude pulmonary mechanisms as a potential cause for heart-voice coupling.

## II. METHOD

### A. Participants

This study has been approved by the Ethics Committee Social Sciences (ECSS) of Radboud University (reference nr.: 22N.002642). The study was originally designed to investigate kinetic effects of arm movements on the respiratory-vocal system (see Pouw et al., 2025; Werner et al., 2024). As planned and supported by a power analysis (see preregistration), we recruited 17 participants: 7 female, 10 male, with an age of 28 ± 6.50 years (M±SD), a body weight of 72.10 ± 10.20 kg (M±SD), a body height of 175.10 ± 8.50 cm (M±SD), a BMI of 23.40 + 2.20 and a triceps skinfold of 19.10 ± 4.30 mm (M±SD). Participants were able-bodied and did not have any constraints in performing the task. All participants provided informed consent before the start of the experiment.

### B. Design

This study was based on a 2 x 2 x 5 experimental design, with the within subject factors of “sound” (vocalization vs. silent expiration), “weight” (weight vs. no-weight) and “movement” (flexion, extension, internal rotation, external rotation, no movement). For a more detailed outline of that experiment, we refer the reader to Pouw et al. (2025). With 4 repetitions per condition, the experiment contained 80 trials in total. For the present study, however, we only used the 16 no-movement conditions. Whether participants were wearing a 1-kg wrist-weight or not, was not of interest, as there were no movements being performed in our target trials. The statistical confirmation of the absence of an effect of wearing a weight on heart-voice coupling can be found in the extended results. Accordingly, we further ignore this weight manipulation.

### C. Procedure and measurements

Before the start of the experiment, we instructed participants about the global aim of the study. Subsequently, we asked participants to take off their shoes, after which we conducted a series of body measurements. We measured their body weight, body height, forearm length, upper arm length, triceps skinfold, and upper arm circumference. We then applied surface electromyography (sEMG) to measure the activity of four muscles in the upper body: the pectoral major, the infraspinatus, the rectus abdominis and the erector spinae. For this we used a wired BrainAmp ExG system (Brain Products GmbH, Munich, Germany) that sampled at 2500 Hz. Before applying the disposable surface electrodes (Kendall 24mm Arbo H124SG), we prepared the skin surface with a scrub gel (NuPrep) followed by cotton ball swipe with alcohol (Podior 70 %). For each muscle we attached three electrodes (active, reference, ground), resulting in 12 electrodes in total. For an overview of the electrode attachments, see **Figure 1**.

**Figure 1.**
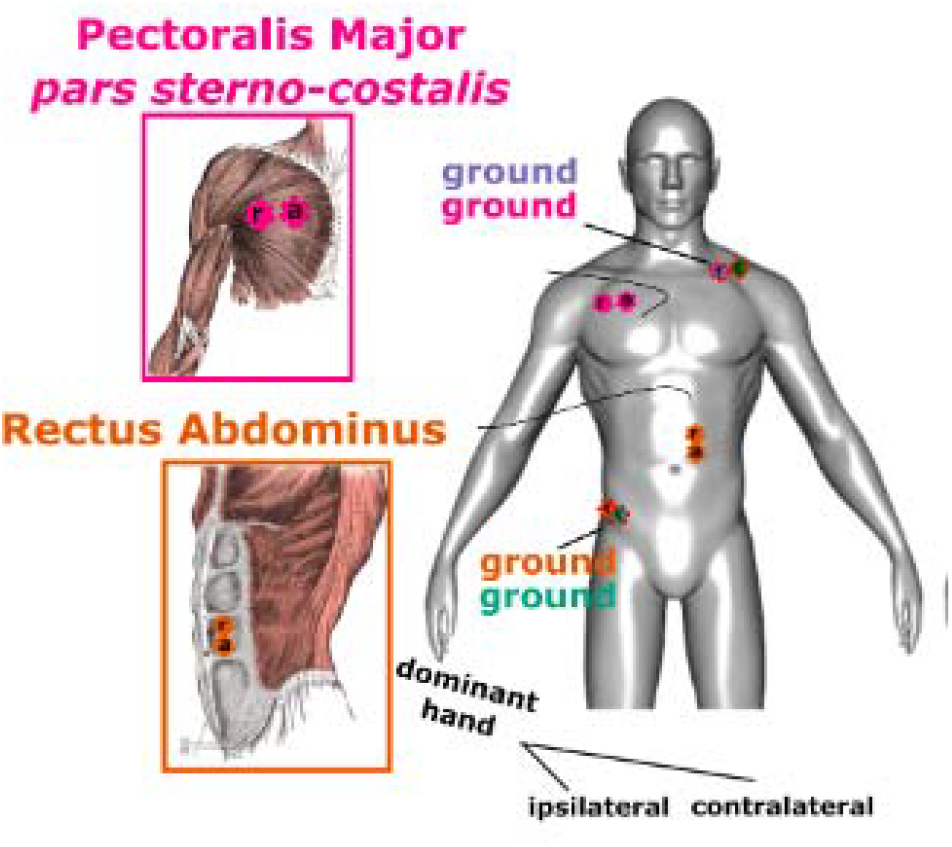
During the experiment we collected surface ElectroMyoGraphy (sEMG) to track muscle activity at four locations: pectoralis major, infraspinatus, which pick up heart signals, but also erector spinae and rectus abdominis (which were not considered here).

Throughout the experiment, we recorded the amplitude of the acoustic signal using a MicroMic C520 (AKG, Inc.) headset condenser cardioid microphone that sampled at 16 kHz. The gain levels of the condenser power source were set by the hardware and were immutable. As part of standard measurements, we instructed participants to breathe silently for 10 seconds without moving their body. Afterwards, participants were given the chance to practice all trial conditions. As explicated, we only discuss no-movement conditions for the current investigation.

Participants were instructed to take a standing position in which they rested their arms alongside their body. For each trial, they were guided by the information on the monitor, which displayed the conditions that were to be performed. In the ‘no movement’ condition (for the other conditions see Pouw et al., 2025) they were instructed to maintain their posture throughout the trial. In vocalization conditions, participants had to produce a sustained vowel (/a/) phonation. After prompting the experimenter that they were ready to start, they were instructed to inhale deeply with a timer counting down for 4 seconds. Then, they had to produce the vowel in a steady-state manner for as long as possible with pursed lips (but not generating a whistle). The procedure for the expiration condition was similar to the vocal condition, only now participants had to expire silently (that is, without producing any sound other than expiratory noises). We interpreted the acoustic noise produced by the silent expiration as the ‘expiration amplitude’. Thus, participants had abducted vocal folds during this condition and we picked up expiratory noises from the head-mounted microphone. In the weight conditions, as opposed to the non-weight conditions, participants wore wrist weights (TurnTuri sports) with a mass of 1 kg. After practicing all conditions, participants performed 80 trials in blocked order, of which 16 were relevant for the present analyses.

### D. Signal preparation

The acoustic and sEMG signals were pre-processed to find signatures of the heartbeat in the amplitude of the voice. This is a deviation from the signal processing in Pouw et al. (2025), where we aimed to *remove* the heart rate from the sEMG. For the sEMG signals, we used a zero-phase 4^th^ order Butterworth filter with a pass-band between 0.75 and 3 Hz to only include the biological range of the heart rate and approximate the standard electrocardiogram (ECG). As expected because of its location close to the heart, the signal derived from the pectoralis major sEMG best reflected the cardiac activity. After filtering, the ECG signals were normalized to z-scores using the mean and standard deviation from the section of the trial from 1 to 6 seconds. Henceforth, we will refer to the filtered and normalized EMG/ECG timeseries of the pectoralis major as the “cardiac activity”. In **Figure 2**, some typical examples of the cardiac activity are shown. To extract the heart rate from the cardiac activity, we applied a Fourier transform to cardiac activity and determined the frequency component with the highest magnitude. This frequency was compared against the frequency we derived from a peak finding algorithm, to test its validity as a heart rate (see extended results notebook). As a last step, we multiplied the frequency by 60 to convert it to beats per minute (bpm), which is more conventional for heart rates.

**Figure 2.**
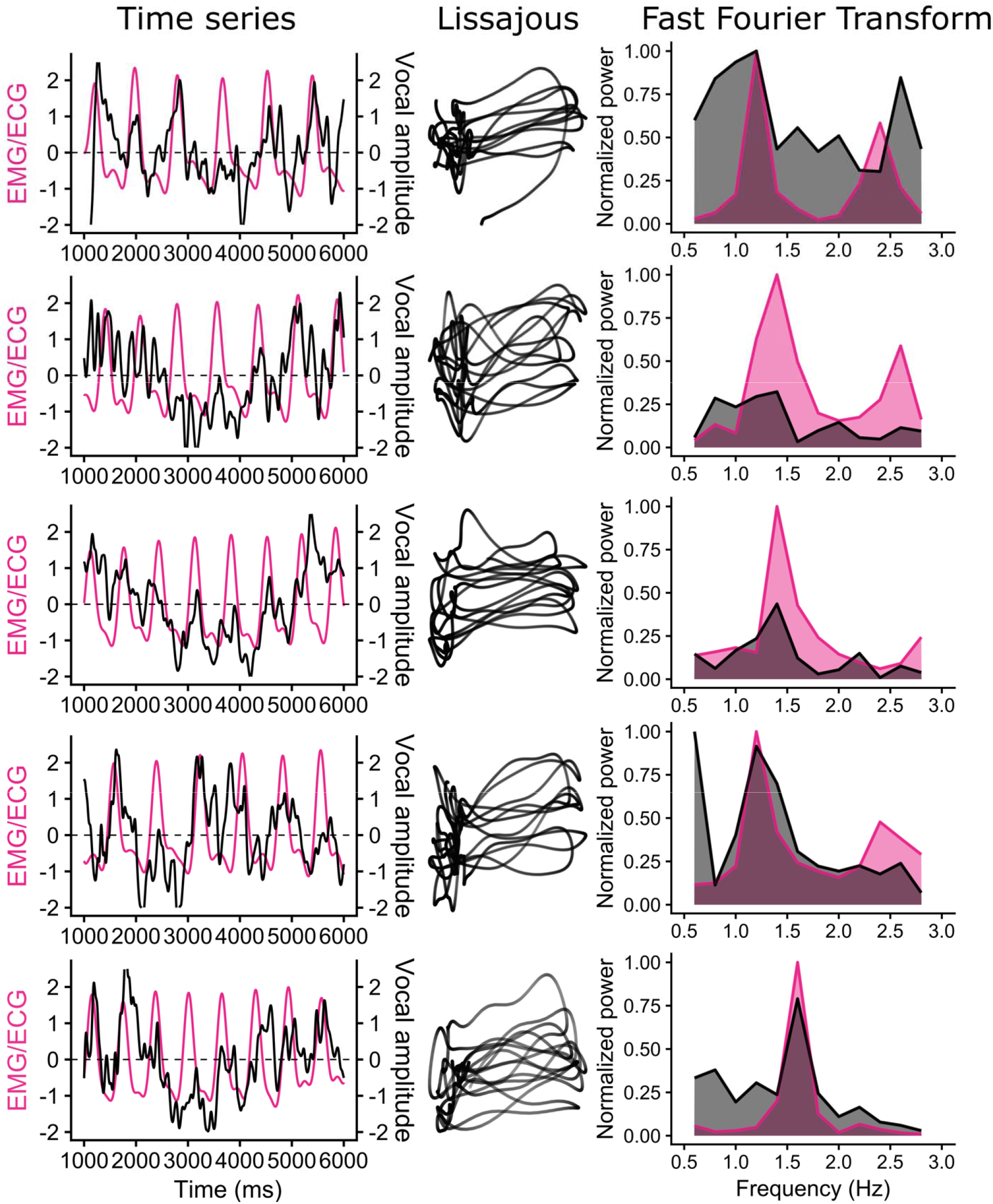
Example of 5 trials of a representative participant (participant 8). Left: the normalized and detrended amplitude envelope timeseries in black and the normalized EMG/ECG timeseries from the pectoralis major in pink. Middle: Lissajous coupling motifs (x = amplitude, y = EMG/ECG). Right: the normalized power spectra (FFT) of the amplitude envelope in black and the EMG/ECG signal in pink.

**Figure 2.**
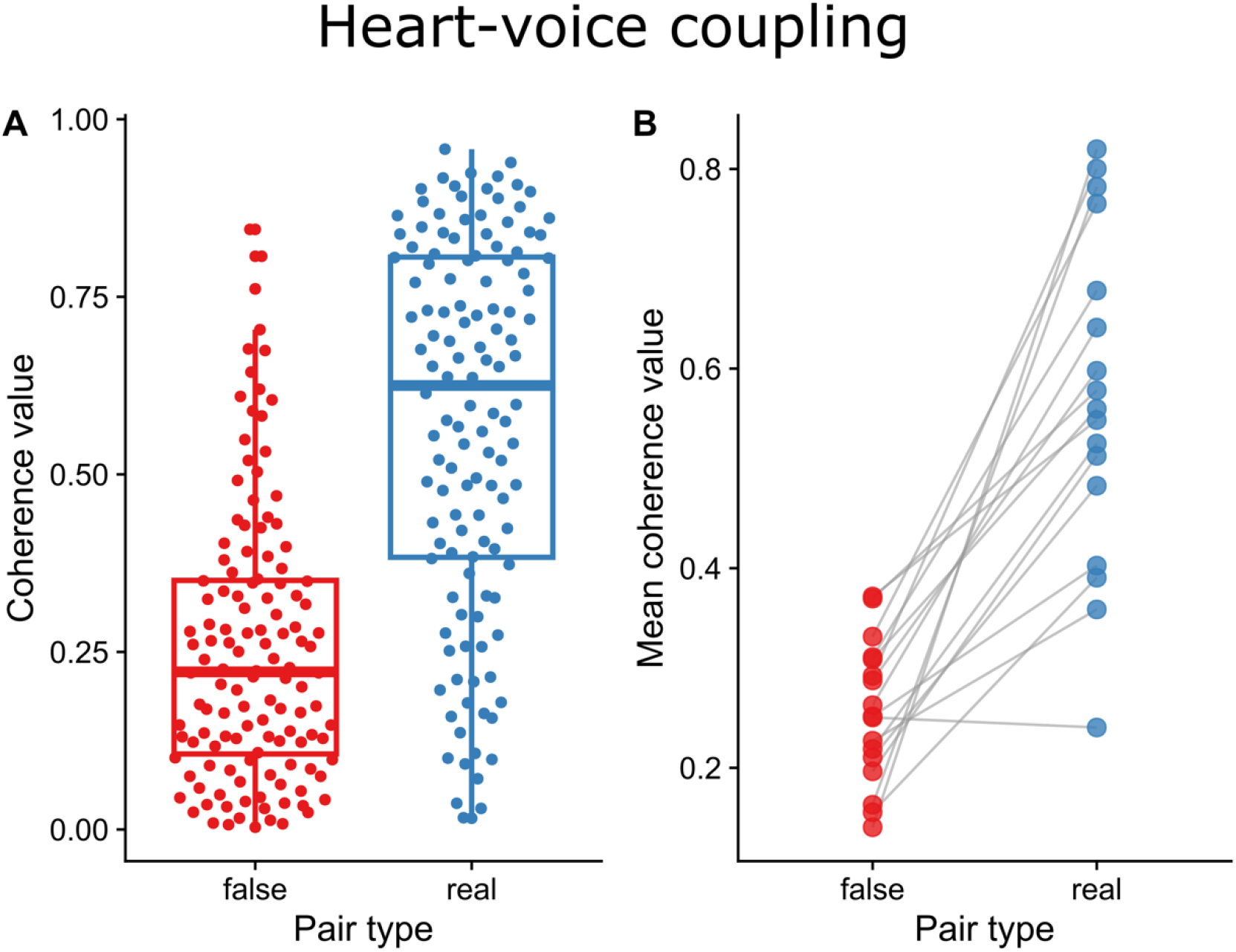
The coherence between the cardiac activity and the acoustic signal in real pairs and false pairs during vowel vocalization, per trial (left) and averaged per participant (right). Coherence values were higher for real pairs than for false pairs, demonstrating that heart-voice coupling was above chance levels.

From the acoustic signal we extracted an envelope that reflected the amplitude of the produced sound (vocalization or silent expiration). We applied a Hilbert transform to the waveform of the acoustic signal, after which we took the complex modulus to create a 1D time series. These timeseries were then smoothed using a Hann filter with a Hanning Window of 12 Hz. We merged the envelopes and EMG signals based on their common timestamps (as provided by the Long-term Spectrum Level; LSL), and then upsampled the envelope using linear interpolation to 2,500 Hz so that we had an equal number of samples compared to the EMG signal. Subsequently, the envelope was normalized to z-scores using the mean and standard deviation from the middle section, i.e. 1 to 6 s. As a last step, we detrended the envelope to remove the gradual decline in amplitude when the lungs reduce in volume and be able to focus on systematic deviations from the trend line. Henceforth, we will refer to this filtered, detrended and normalized amplitude envelope signal as the “acoustic amplitude” in short. In **Figure 2**, some typical examples of the acoustic amplitude relative to the heart rate signals are shown. The figure shows the relationship in the temporal domain (timeseries, left), the spatial domain (Lissajous coupling motives with x = amplitude and y = ECG, middle) and the frequency domain (power spectra, right)

Using the cardiac activity and the acoustic signal as base signals for all further analyses, we conducted a series of tests to find evidence for heart-voice coupling and to discriminate between the vocal fold and pulmonary mechanisms behind the heart-voice coupling.

### E. Hypothesis testing

As a first step, we investigated whether the acoustic signal was coherent with the cardiac activity. More specifically, we computed the squared-magnitude coherence between the amplitude envelope and the ECG signal at the cardiac frequency, as extracted earlier with the Fourier analysis. Note that the squared-magnitude coherence expects a consistent relative phase between the signals (regardless of the value of the phase relationship). If the coherence values at the cardiac frequency would be larger than chance, this would confirm heart-voice coupling. Accordingly, we not only computed the coherence between real pairs of cardiac activity and acoustic signal, but also between false pairs (surrogate analysis). These false pairs were created by randomly combining the timeseries across trials, making sure that no combination of cardiac activity and acoustic amplitude occurred twice. A significantly higher coherence for real pairs than for false pairs would then indicate that the coherence between the timeseries is above chance. To test this statistically, we assessed whether a mixed linear regression model that included pairing type (real pairs vs. false pairs) would explain the variance in the coherence better than a base model that only predicted the overall mean.

Secondly, we tested whether heart-voice coupling would be more pronounced at the beginning of the vocalization –when the lungs were full of air– than at the end. We compared the squared-magnitude coherence between the acoustic signal and the ECG signal in the first half of the vocalization (from 1001 to 3500 ms), to the coherence in the second half (from 3500 to 5999 ms) for real and false pairs. As a statistical test, we included pairing type (real pairs vs. false pairs) and trial half (first half vs. second half) in a mixed linear regression model to see if it explained the variance in the coherence better than a base model that only predicted the overall mean.

Thirdly, we tested whether the amplitude of a silent expiration, like that of a sustained vocalization, would also be coherent with the cardiac activity. If so, this would provide evidence in favour of pulmonary mechanisms behind heart-voice coupling, because the vocal folds are abducted and thus not involved in expiration. For the expiration conditions, we computed the squared-magnitude coherence between the acoustic amplitude and the cardiac activity, again for false as well as random pairs. If the mixed linear regression model that included pairing type (real pairs vs. false pairs) explained the variance in the coherence better than a base model that only predicted the overall mean, we concluded that the coherence between expiration amplitude and cardiac activity was above chance level.

For all analyses the significance level was set to α = 0.05. Where appropriate we reported mean, standard deviation and 95% confidence intervals as M ± SD, [CI_min,_ CI_max_].

### F. Exploratory analyses

We also conducted some exploratory analyses. First, we investigated whether heart-voice coupling was dependent on the heart rate, speculating that the impact of the heartbeat on the acoustic amplitude may be more pronounced when the heart beats faster. To do so, we included the extracted heart rate as a factor in our mixed linear regression model and assessed whether this model better predicted the variance in the coherence values than a base model. Second, we investigated whether the onset of the vocalization was coordinated with the beating of the heart, to explore whether the heart beat may affect the timing of speech. We defined vocalization onset as the first peak in the velocity (attack phase) of the amplitude and computed the time interval between this onset and the closest peak in the cardiac activity. Subsequently, we normalized this time distance to the duration of the cardiac cycle and transformed it to degrees to acquire the relative phase value. If the onset of the vocalization were to be coordinated with the heartbeat, we would see the emergence of a stable relative phase between the two events. To assess whether the relative phase distribution was significantly different from chance, we again compared real pairs to false pairs.

## III. RESULTS

All analysis code and extended results are available in quarto/RMarkdown notebook format. There is an extended result notebook for heart-voice coupling (https://wimpouw.github.io/VoiceAndHeart/) and the heart-expiration coupling (https://wimpouw.github.io/VoiceAndHeart/heartrate_expire).

### A. Amplitude of vowel vocalization is coherent with cardiac activity

In **Figure 3**, we plotted the coherence between acoustic amplitude and cardiac activity during vowel vocalization for all trials of all participants. The boxplot on the left shows the coherence values for false pairs, whereas the boxplot on the right shows the coherence for real pairs, with the middle line indicating the median coherence per group. Comparison of the boxplots reveals that the coherence values are higher for real pairs (0.572 ± 0.264, 95%CI [0.526, 0.618]) than for false pairs (0.256 ± 0.193, 95%CI [0.222, 0.289]). This was statistically confirmed by a mixed linear regression model that included pairing type (real pairs vs. false pairs) as a main factor, and trial nested in participant as random intercept. This model better explained the variance in the coherence than a base model that predicted the overall mean (Change in *X*^2^ (1) = 106.45, *p* < .001). All model coefficients are shown in **Table I**.

**TABLE I.**
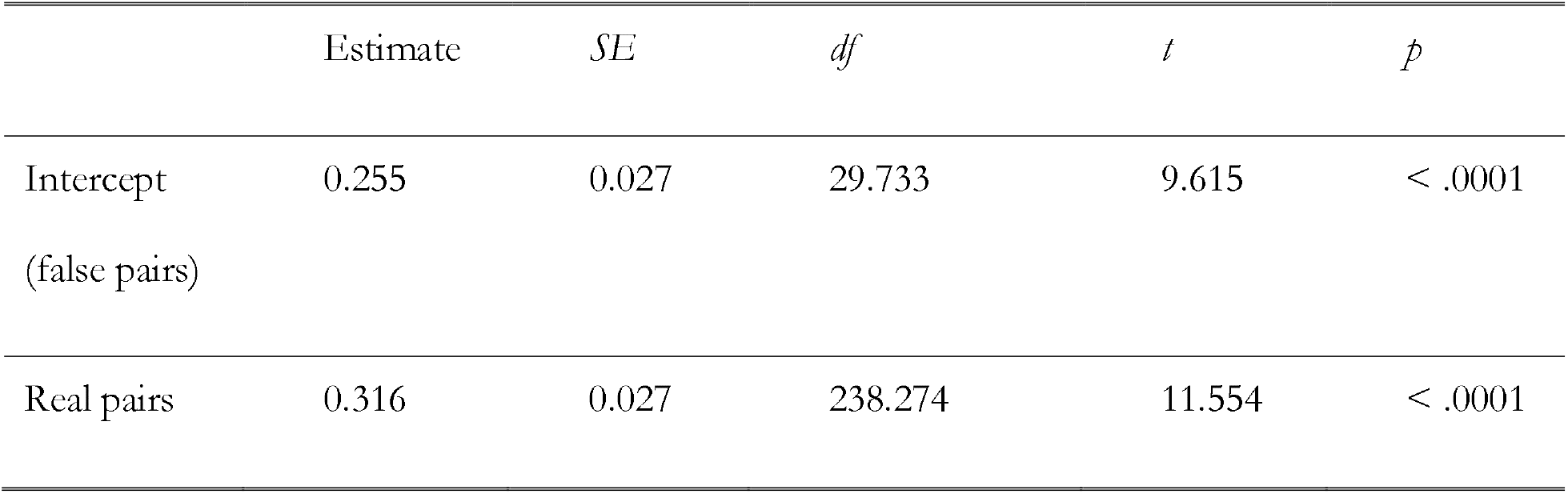
Effects of pairing type on coherence values in mixed linear regression model.

**Figure 3.**
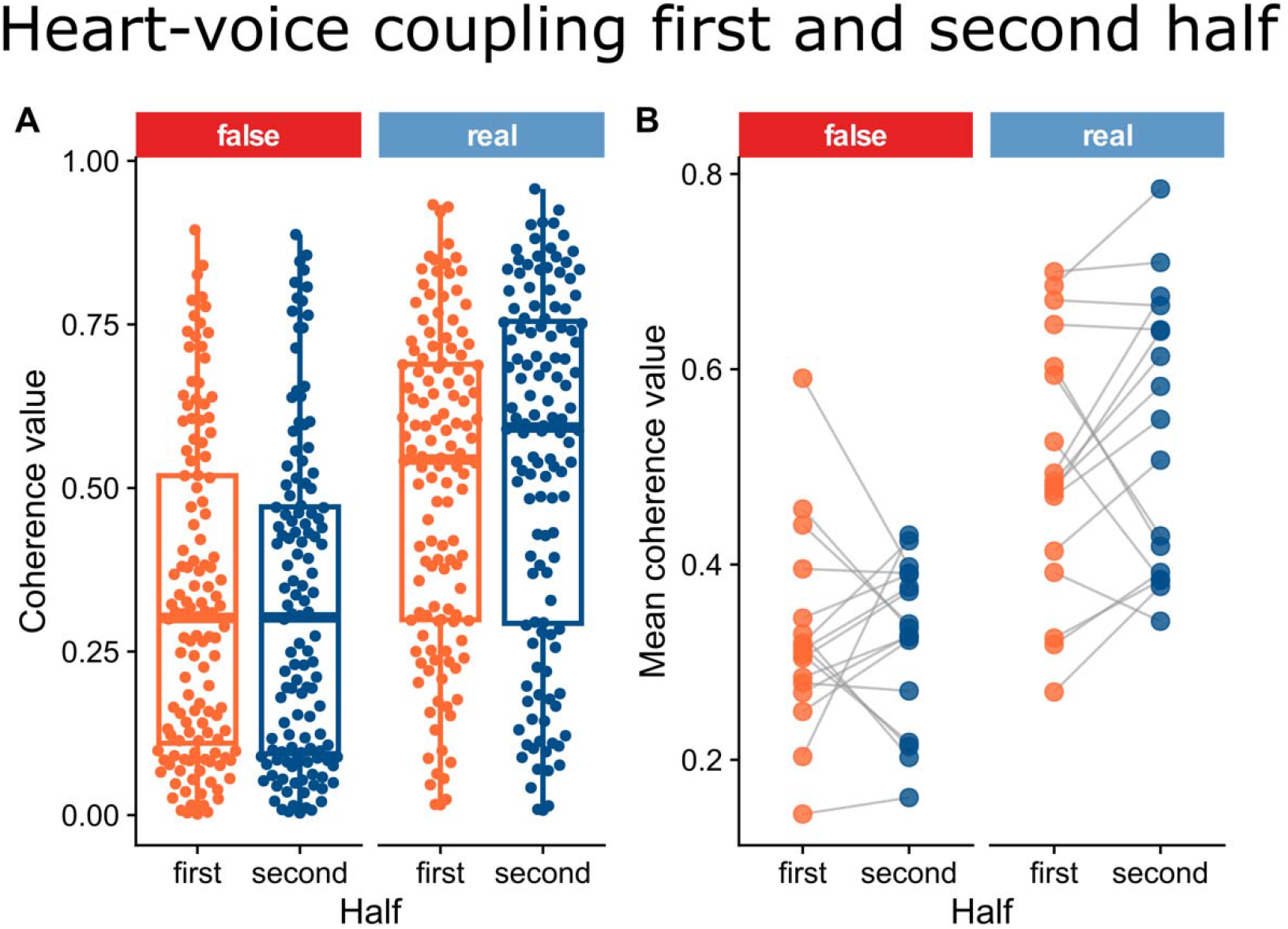
The coherence between cardiac activity and acoustic signal in real pairs and false pairs, compared between the first and the second half of the vowel vocalization. The graph on the left shows the coherence values per trial, whereas the graph on the right shows the coherence values averaged per participant. There was no significant difference between the first and the second half of the vocalization and no interaction effect.

### B. Heart-voice coupling does not change over the course of the vocalization

In **Figure 4** we plotted the coherence between acoustic amplitude and cardiac activity in the first and second half of the vowel vocalization, for all trials of all participants. The boxplots on the left show the coherence values for false pairs, whereas the boxplots in the right show the coherence values for real pairs, with the middle line indicating the median coherence per group. Comparison of the boxplots reveals that the coherence values were not drastically different between the first half (false pairs: 0.326 ± 0.242, 95%CI [0.284, 0.368], real pairs: 0.503 ± 0.240, 95%CI [0.461, 0.545]) and the second half (false pairs: 0.321 ± 0.244, 95%CI [0.278, 0.364], real pairs: 0.537 ± 0.268, 95%CI [0.490, 0.584]) of the vowel vocalization. This was statistically confirmed by our mixed linear regression model with participant as random intercept. Whereas the inclusion of pairing type (real pairs vs. false pairs) and half (first half vs second half) as main effects improved the model fit (change in *X*^2^ (2) = 94.505, *p* < .001) the inclusion of a pairing type x half interaction effect did *not* do so (change in *X*^2^ (1) = 1.043, *p* = 307). All model coefficients are shown in **Table II**.

**TABLE II.** Effects of pairing type and vocalization half on coherence values in mixed linear.

|  | Estimate | *SE* | *df* | *t* | *p* |
| --- | --- | --- | --- | --- | --- |
| Intercept (false half, false pairs) | 0.326 | 0.026 | 44.617 | 12.537 | < .001 |
| Second half | -0.005 | 0.027 | 381 | -0.187 | 0.852 |
| Real pairs | 0.177 | 0.027 | 381 | 6.562 | < .001 |
| Second half x<br>real pairs | 0.039 | 0.038 | 381 | 1.018 | 0.309 |

**Figure 4.**
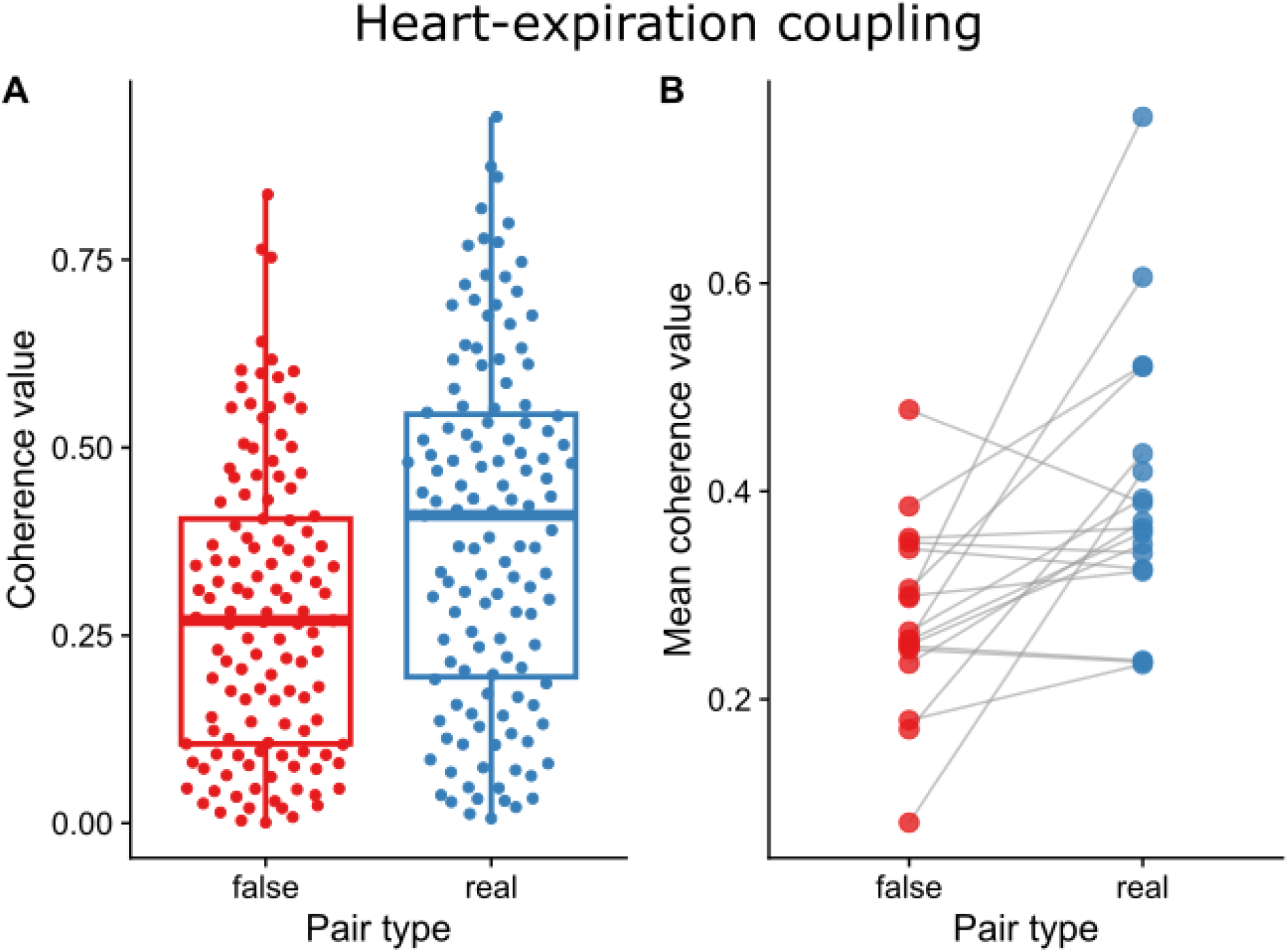
Coherence between cardiac activity and acoustic signal in real and false pairs during silent expiration. Coherence values are higher for real pairs than for false pairs, demonstrating that heart-voice coupling during silent expiration without speech was above chance level (5%).

Please note that we also performed a corroborating analysis that conserved a better time-frequency resolution. Namely, we applied a cross-wavelet analysis where we extracted the time-dependent cross-wavelet coherence. The results were similar to those reported here (for further details see the extended results).

### C. Coherence is also observed between expiration amplitude and cardiac activity

In **Figure 5**, we plotted the coherence between acoustic amplitude and cardiac activity during silent expiration (without speech) for all trials and all participants. The boxplot on the left shows the coherence values for false pairs, whereas the boxplot on the right shows the coherence for real pairs, with the middle line indicating the median coherence per group. Comparison of the boxplots reveals that the coherence values are significantly higher for real pairs (0.389 ± 0.231, 95%CI [0.348, 0.430]) than for false pairs (0.281 ± 0.192, 95%CI [0.247, 0.315]), showing that the coherence between acoustic signal and cardiac activity during silent expiration was above chance levels. This shows that the heartbeat cannot only be extracted from a vowel vocalization, but also from the acoustic noise of an expiration without speech. The result was confirmed by the mixed linear regression model that included pairing type (real pairs vs. false pairs) as a predictive factor. The model better explained the variance in the coherence than a base model that predicted the overall mean (change in *X*^2^ (1.00) = 17.091, *p* < .001). All model coefficients are shown in **Table III**.

**TABLE III.** Effects of pairing type on coherence values in mixed linear regression model.

|  | Estimate | *SE* | *df* | *t* | *p* |
| --- | --- | --- | --- | --- | --- |
| Intercept (false pairs) | 0.283 | 0.023 | 33.046 | 12.084 | < .0001 |
| Real pairs | 0.108 | 0.026 | 232.555 | 4.203 | < .0001 |

### D. Exploratory analyses

The coherence between the acoustic amplitude and the cardiac activity during vowel vocalization was not dependent on the heart rate of participants. Also, the onset of vocalization was not temporally aligned with the beating of the heart. Note further that all analyses for the vocalization condition we presented earlier were also replicated for expiration trial conditions. For an overview of these analyses, see the extended results.

## IV. DISCUSSION

The existence of heart-voice coupling has been known for several decades (Orlikoff & Baken, 1989a) and technological advances have been made to extract the heartrate from the voice, with many potential applications (Mesleh et al., 2012; Sakai, 2015; James, 2015; Smith et al. 2017; Milton & Monsely, 2018). Notwithstanding, the physiological origin of the coupling between heart and voice has been left unexplained. This is unfortunate, because the quality of algorithms that extract HR from vocal acoustics is ultimately dependent on our mechanistic understanding of heart-voice coupling. In the current contribution, we built upon the seminal work of (Orlikoff & Baken, 1989a) and replicated their finding of heart-voice coupling. The acoustic amplitude of a sustained vowel (/a/) vocalization was found to be significantly coherent with cardiac activity, confirming a coupling of the cardiovascular system to the vocal system. But does the heartbeat enter the voice via the vascularization of the vocal folds, or via a subglottal interaction between the beating heart and the richly perfused lungs? To discriminate vocal fold from pulmonary mechanisms and to provide a physiological explanation for heart-voice coupling, we conducted a series of empirical tests.

First, we investigated whether the coupling between heart and voice was dependent on the inflation of the lungs. If the coupling were to be more pronounced at the beginning of the vocalization –when the lungs are inflated– than at the end, this would suggest a mechanical interaction between the heart and the lungs, providing support for pulmonary mechanisms. Our data showed, however, that this was not the case. Between the first and the second half of the vocalization, there was no significant difference in the coherence of the acoustic amplitude with the cardiac activity. This result can indicate two things. On the one hand, it is possible that there is no subglottal interaction between the heart and the lungs. The mechanical coupling of the heart to the lungs, via the pericardium, may be diminished by the elastic properties of the thorax. The thorax follows the lungs’ deflation to great extent and encapsulates the entire dynamic recruitment of other stabilizing actions. Indeed, there is evidence that humans can stabilize pulmonary pressures by recruiting additional respiratory muscle actions when elastic recoil diminishes their expiratory drive (Bouhuys et al., 1966; Ohala, 1990). On the other hand, our finding may indicate that the interaction between the heart and the lungs is too small for it to be identified in the acoustic signal. In any case, the test employed was not conclusive. Therefore, we conducted a second empirical test to find evidence in favour of (or against) the pulmonary mechanisms: we assessed the acoustic signal of a silent expiration.

During expiration, the vocal folds are abducted, so they cannot contribute to expiratory modulation. Nonetheless, we found a significant (i.e. above chance) coherence between the acoustic signal and the cardiac activity during conditions of silent expiration. This means that the heart’s activity is manifested in the acoustic signal, *even when the vocal folds are not involved in its production*. Accordingly, it refutes the widely held –but never directly tested– assumption that heart-voice coupling is only caused by vocal fold mechanisms (Orlikoff, 1989; Sakai 2015; Poleshenkov & Basov, 2020). Instead, it suggests that the heartbeat can also manifest itself in the voice via an interaction between the heart and the lungs that drive fluctuations in the pulmonary pressure. After all, the subglottal pressure is the prime determinant of the acoustic noise during a silent expiration. The interaction between the heart and the lungs may take two forms. On the one hand, the pulmonary pressure may fluctuate because the blood pressure in the aorta, pulmonary arteries and lung vessels rises and falls with the beating of the heart. This has been proposed by Poleshenkov & Basov (2020) and would explain why signatures of the heartbeat were also found in the expiration amplitude here. On the other hand, the heart may have a mechanical impact on the lungs, to which it is connected via the pericardium. This, too, could cause fluctuations in the flow of air from the lungs and thus in the pulmonary pressure. Future research into the dynamics of the heart-lung interaction could discriminate between these pulmonary mechanisms.

### A. The heart as a pattern generator for speech

Taken together, our results suggest that heart-voice coupling is a replicable phenomenon with multiple physiological causes. From the perspective of speech production, this means that the vocal system is confronted with rhythmic perturbations from the cardiovascular system that may or may not be accounted for. As such, heart-voice coupling is not only relevant for the development of medical technology, but also for the study of psycholinguistics. In the extreme, human speech may have evolved by being scaffolded on weakly coupled pacemakers such as the beating of the heart. Minimally, the heart may be a nuisance to voice control that contributes to fluctuations in the voice quality. In any case, the heart adds a fundamental rhythm to the multitude of already appreciated hierarchically nested rhythms of which human speech is composed (Kelso & Tuller, 1984; Saltzman & Munhall, 1989; Tilsen & Arvaniti, 2013; Poeppel & Assaneo, 2020). For instance, speech involves vocalization embedded in labial and tongue movements, which are embedded in jaw cycles (MacNeilage, 2010), which in turn are embedded in the chest movements (Rochet-Capellan & Fuchs, 2013). Moreover, recent studies have shown that the voice is affected by physically perturbing action such as gestures (Pouw & Fuchs, 2022; Pouw et al., 2025). Heart-voice coupling thus complexifies our understanding of the rhythmic underpinning of speech, because it shows that the vocal system is even affected by the most primordial, life-preserving pattern generator in the body: the heart.

The conceptualization of speech as an integration of rhythms puts the heart-voice coupling into a functional perspective. It raises questions of how the heartbeat may affect vocal control or the timing of speech. It is well known, for example, that the weakly coupled perturbation of locomotor strides onto the respiratory system can lead to a synchronization of respiratory cycles with locomotion in (human) animals (Pouw & Fuchs, 2022). Could the same be true for the cardiac cycle? Providing sufficient answers to such questions is beyond the scope of this paper. However, we did perform some explorative analyses that allow us to speculate, briefly, on this matter.

We explored whether the onset of the vocalization is synchronized to the beating of the heart, which was not the case. The reason for this could be that the task was not unconstrained enough, in the sense that the cue for participants to initiate the vocalization was dictated by the experiment itself and therefore too strong. Alternatively, it is possible that the cardiac cycle only plays a marginal role in the coordination of speech or that its role is more complex, for instance by tying in with the respiratory cycle.

Other questions concern whether the information of heart beats in vocal patterns can serve as a basis for physiological synchrony (Wicher et al., 2025), as a source of bodily meaning for sound-symbolism (Ćwiek et al., 2025), or as an explanatory basis for where vocal isochrony (equal timing between vocal sounds) comes from in evolution in humans and other animals (Burchardt et al., 2019; Larsson et al., 2019; Ravignani et al., 2019). Furthermore, we wonder whether people can perceive the heart rate fluctuations by closely attending to vocal sounds (Pouw et al., 2020; Ghazanfar et al., 2007; Pisanski et al., 2016), or whether humans and other animals are implicitly influenced by the vocal features the heart modulates in the perception of the voice (Anikin, 2023; Rathcke et al., 2024; Valente et al., 2025), such as emotions which are related to vocal acoustics (Schewski et al., 2025; Yao et al., 2012). Our findings underline that there is an acoustic medium that can carry heart rate, which may serve as the start of an explanatory multimodal account of how one can informationally ground between-person heart rate synchronization (Coutinho et al., 2021; Wicher et al., 2025).

While the current data and experiment are not sufficiently rich to answer these questions, we hope our results revive interest into heart-voice coupling and its functional implications.

### B. Conclusion

Within behavioral and neurosciences, it is becoming increasingly clear that behavior requires the capacity to maintain stable endogenous rhythms (Burchardt et al., 2019; Ghazanfar, 2013; Hamilton et al., 2025; Kelso & Tuller, 1984; Larsson et al., 2019; MacNeilage, 2010; Pouw et al., 2021; Rathcke et al., 2021; Ravignani et al., 2019; Wilson & Wilson, 2005). We are not only responsive to the exogenous flux of information from the environment. Our behaviour also depends on the physiological cycles within our bodies, such as the rhythm of the heart (Duschek et al., 2013). Rhythmic finger tapping is timed in coordination with the heartbeat (Davidson et al., 1981) and the initiation of action is timed to the phase of the cardiac cycle (Saari & Pappas, 1976; Jennings & Wood, 1977; Rae et al., 2018; Kimura et al., 2023), to give some examples. In the current contribution we showe d that vocal acoustics, too, reflect signatures of cardiac activity. We demonstrated that both vocalization and silent expiration amplitudes contain signatures of the heartbeat, suggesting that heart-voice coupling must at least have pulmonary origins. By invoking functional questions and potential avenues for further research, we hope that our study will bring new life to investigations of heart-voice coupling. Ultimately, this will produce useful applications in medical, psychological and applied fields, as well as a deeper understanding of the physiology underlying human speech.

## V. ACKNOWLEDGEMENTS

We would like to thank the participants who volunteered in this study. We would like to thank Pascal de Water for his technical support in setting up the multimodal LSL-synchronized lab.

## VII. AUTHOR DECLARATIONS

### B. Ethical approval

This study has been approved by the Ethics Committee Social Sciences (ECSS) of Radboud University (reference nr.: 22N.002642).

## VIII. DATA AVAILABILITY STATEMENT

Extended results and all codes for analysis are available in quarto/RMarkdown notebook format. There is an extended result notebook for heart-voice coupling (https://wimpouw.github.io/VoiceAndHeart/) and the heart-expiration coupling (https://wimpouw.github.io/VoiceAndHeart/heartrate_expire). The full dataset (64) with masked videos to reduce identity exposure is openly available on the Radboud Data Repository and forms the input for the scripts provided: https://doi.org/10.34973/p9se-mq71

## IX. FOOTNOTES

1. In contrast with the original publications of Orlikoff and Baken (1989), these studies did not extract the full phase progression of the cardiac cycle from the acoustic signal. Instead, they only predicted the average HR from the voice recordings and compared the predicted to the measured HR using correlation measures.
2. The subglottal pressure is strongly related to the alveolar pressure, i.e., the air pressure in the lungs. For simplicity, we will not consider the subtle difference between the two here.
3. Known as the cardioballistic effect (Orlikoff 1989b).
4. Orlikoff (1990a) has mentioned performing a similar empirical test, thereby failing to find a significant effect. However, he did not report the methodology underlying his test, nor did he present concrete results to support his conclusion.

